# Coupled Biomechanical and Ionic Excitability in Developing Neural Cell Networks

**DOI:** 10.1101/2023.02.15.528510

**Authors:** Sylvester J. Gates, Phillip Alvarez, Kan Cao, Kate O’Neill, Wolfgang Losert

## Abstract

Waves and oscillations play a key role in the flow and processing of information in the brain. Recent work has demonstrated that in addition to electrical activity, biomechanical signaling can also be excitable and thus capable of self-sustaining oscillations and waves. Here we measured the biomechanical dynamics of actin polymerization in neural precursor cells throughout their differentiation into populations of neurons and astrocytes. Fluorescence-based live-cell imaging allowed us to analyze the dynamics of actin in conjunction with the dynamics of calcium signals. Actin dynamics throughout differentiation showed a rhythmic character, localized mostly in processes, with changes in scale associated with differentiation. Furthermore, actin dynamics impact ionic dynamics, with an increase in the frequency of calcium bursts accompanied by a decrease in cell-cell correlations when actin dynamics is inhibited. This impact of cytoskeletal dynamics on cell-cell coupling and ionic neural cell signaling suggests that information flow in the brain may be able to harness both biomechanical and electrical/ionic excitability.

## Introduction

The development of neural networks is an amazing feat of nature that results in the emergence of collective ionic activity, and in much of the animal kingdom, these rhythms ultimately support the emergence of thinking and consciousness. Much of the literature on neural networks is neuron-centric in focus and measures *electrical* excitability within *neurons*. This view of neural network excitability misses two key aspects of the brain: (1) the development of the brain involves a variety of cell types, the most common among them astrocytes that act as modulators of neuronal cells, and (2) neural network development also encompasses biophysical morphological changes of neural cells and networks, raising the question of the role of biomechanics in the emergence and sustainment of collective ionic activity ^1–6^.

One starting point for neural network development are neural precursor cells (NPC), a mixed population of cells containing both neural stem cells and neural progenitor cells, which differentiate downstream into populations of mature neurons, astrocytes, and oligodendrocytes within the brain^7^. During differentiation, profound changes occur within each cell as the cytoskeleton rearranges from immature NPCs to the differentiated mostly post-mitotic state that characterizes more mature neural cells^1,8–10^. At the same time, cell groups develop communications that characterize networks of mature neural cells.

Neurons differentiated from NPCs are the canonically “electrically active” cells associated with rapid changes in transmembrane potential^11–13^. Neuronal action potentials (AP) are tightly regulated through the influx and efflux of sodium and potassium ions that regulate the transmembrane potential of the cells rapidly on timescales of milliseconds. Other ions also transit across the membrane during the AP including Ca^2+^, which moves out of the cell in tens to hundreds of milliseconds making it a useful tool for monitoring neuronal activity in real-time without the need for millisecond timescale imaging ^14–16^. Several technologies take advantage of these slower Ca^2+^ transients, namely calcium-sensitive dyes (like Fluo-4, X-Rhod, and CalBryte) and genetically encoded calcium indicators (like the GCaMP family of proteins)^17,18^.

Astrocytes, which are also differentiated from NPCs, perform a variety of tasks such as maintaining the cells that form the blood-brain barrier, releasing gliotransmitters in response to calcium and neuronal signaling, and forming scar tissue in response to infection and damage within the Central Nervous System ^1^. Healthy astrocytes have a star-like morphology, with the star’s tips thought to modulate and interact with the synapses of neurons ^1–3^. While astrocytes unlike neurons do not depolarize, recent studies have shown that they are not “silent” and instead show oscillations and waves in intracellular calcium levels, indicating an excitable character of astrocytic Ca^2+^ signaling ^2,19^. Network-wide Ca^2+^ signaling in NPCs can form the basis for communication in more mature neural networks ^20^. This Ca^2+^ signaling in astrocytes can be modulated by some of the same physiologically relevant inputs that also alter neuronal activity such as ATP, glutamate, and noradrenaline ^3,19,21^. Glial cells, often overlooked in traditional neuron-centric views, play a pivotal role in modulating neuronal networks^22^. Through their diverse functions, including neurotransmitter uptake, ion homeostasis, and synaptic pruning, glial cells actively contribute to shaping neural circuits and information processing^1,2,23–25^. Astrocytes, for instance, regulate extracellular ion concentrations and neurotransmitter levels, impacting synaptic strength and plasticity. Microglia, the immune cells of the brain, engage in synaptic remodeling and influence neuronal connectivity. Understanding the dynamic interplay between glial cells and neurons is essential for unraveling the intricate mechanisms underlying neural network function and dysfunction.

The development of an active neural network also involves cellular biomechanical machinery. Driving much of neural morphological change is actin cytoskeletal polymerization and depolymerization ^26–28^. Mature neurons are not generally migratory; however, they still exhibit intracellular actin dynamics^29–32^. Previous research on other cell types had shown that actin dynamics can be driven by Ca^2+^ dynamics in several contexts including calcium-dependent actin reorganization in chondrocytes, receptor-induced calcium mobilization mediating actin rearrangement in B lymphocytes, and increases in intracellular calcium leading to actin reorganization and migration in breast cancer cells in response to propofol ^33–35^. Within a neural context a number of studies have shown typically localized coupling between actin dynamics and calcium signaling in dendritic spines – with calcium influx regulating dendritic spine plasticity ^36,37^.

However, recent studies have discovered that the dynamics of the actin cytoskeleton do not necessarily require a specific signaling input, but that the biomechanical actin polymerization machinery acts as an excitable system capable of generating waves or oscillations ^2,38–42^. The key characteristics of biomechanical excitability are the same as for electrical excitability: (1) the system has multiple distinct states, (2) there is some threshold for activation, and (3) there is a refractory period following activation in which another activation is suppressed. Moreover, these recent studies have even suggested that biomechanics and mechanical cues can play a key role in neuronal communication such that mechanical cues like pressure can lead to dynamic structural changes of actin in synaptic spines that can leave longer-lasting (tens of seconds) impacts on the neurotransmission of glutamate and therefore neuronal communication ^29^.

Since actin dynamics exhibit excitability, it opens the possibility that biomechanical waves and oscillations may contribute to the information in neural networks in their own right. In neuronal cells, the dynamics of actin play a role in multiple functionalities: In axons, dynamic actin is found in growth cones during axonogenesis, and in small actin patches and waves, which can propagate perpendicular to the axon. In dendrites, dynamic actin-rich structures can form along the length of the dendrite, and evolve into dendritic spines, important structures for neuronal synaptic communication ^43^. Recent work has demonstrated that the dynamic character of actin remains important in synapses, with drugs affecting actin dynamics modulating and preventing important synapse functions like long-term potentiation ^44,45^. In terms of network populations, recent work has shown co-cultures with neurons and astrocytes, show an increase in actin dynamics specifically in subcellular local hotspots which may suggest these two cells might impact each other’s biomechanical systems during communication ^46^.

Therefore, the goal of our study is to characterize the changes in actin dynamics during neural cell differentiation and elucidate the coupling between actin dynamics and Ca^2+^ activity.

## Results

To investigate actin dynamics and Ca^2+^ in neural cells, we use live optical imaging of immortalized NPCs with genetically labeled actin probes and calcium fluorescent indicators throughout their differentiation ^47^.

### Controlled differentiation in neural progenitor cells with stable lifeactTagGFP2 expression

We established an NPC line with a commonly used genetic actin probe, LifeactTagGFP2, using a commercially available hNPC line immortalized through V-MYC transduction. LifeAct is a small peptide sequence that binds to actin similar to other actin-binding proteins often used in live-cell imaging to track F-actin structures^48^. The hNPC line used shows immature neural stem cell markers nestin and sox2 in the undifferentiated state as assayed by immunohistochemistry (Fig 1A). Differentiation initiated by the removal of growth factors leads to a mixed population of differentiated cells with mutually exclusive expression of beta-3-tubulin (TUBB3) or glial fibrillary acidic protein (GFAP) which are neuron and astrocyte expression markers respectively (Fig. 1B). Lentiviral transduction was used to obtain stable expression of LifeactTagGFP2 within the undifferentiated cells and this expression is retained throughout the induction of differentiation. During differentiation cell morphology of hNPC change dramatically. Undifferentiated cells show polygonal morphology characteristic of many adherent epithelial cell lines. Differentiated cells show a reduction in cell body size, as well as long processes extending tens of hundreds of microns in length (Fig.1). Differentiated populations achieve around 25% (24.1881 ± 9.6076) TUJ1 positive neurons (n = 8 with at least 7 FOV looked at in each sample).

**Figure 1.**
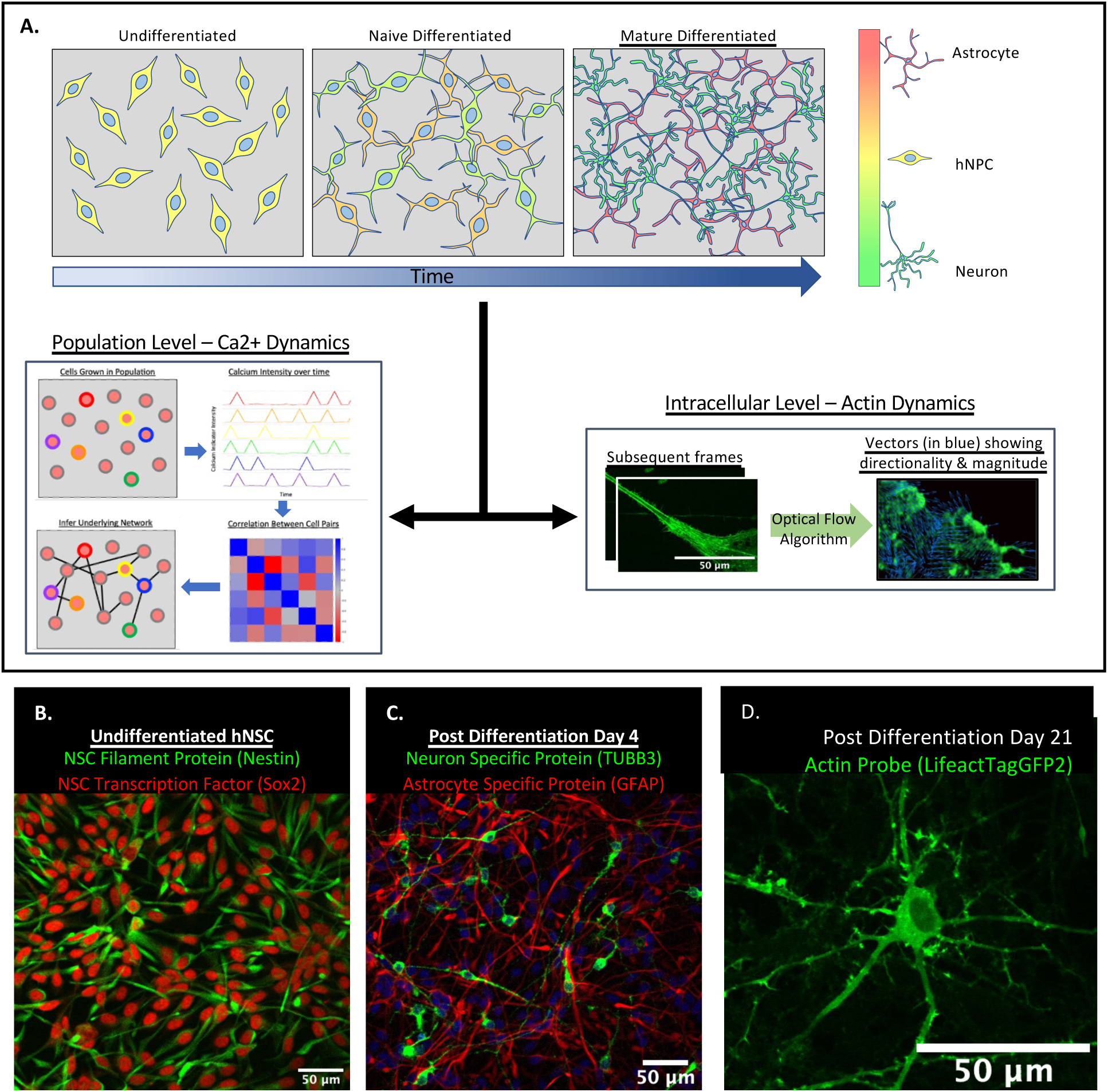
Model hNPC Line Workflow and Differentiation Capability. Workflow for the proposed work (A) starting with timecourse of hNPC differentiating towards matured neurons and astrocytes followed by analysis of calcium and actin dynamics. Representative immunofluorescence image of undifferentiated hNPC (**B**) with expression of Nestin (green) and Sox2 (red). Immunofluorescence on 4 days post differentiation initiation of hNPC (**C**) with expression of TUBB3 (green) or GFAP (red) with nuclear staining (blue). Representative confocal fluorescent image of lifeact-GFP stable transduction of a 21-day post differentiation initiation cell derived from hNSC (**D**).

### Actin dynamics of hNPC have characteristic rhythm and change scale throughout differentiation but not directionality

We analyzed actin dynamics of hNPC throughout differentiation by looking at 3 specific time periods during their maturation: (1) during the undifferentiated cell state, (2) during the early/naive differentiated cell state (3-7 days post differentiation), and (3) during the late differentiated cell state (14-28 days post differentiation). Qualitatively we observed a change in the scale of the actin dynamics during these three stages of differentiation. To observe differences temporally images were constructed representing temporal max-projections – where each frame is represented by a different color and is collapsed into a single image. Areas that show white suggest static regions, while areas of color show dynamics during specific time points (Figure 2). Thus, colored regions can therefore be influenced by both actin dynamics parallel to the cell boundary as well as actin dynamics that changes the cell shape itself, however, these cells are not known to move or change shape rapidly, so actin dynamics independent of cell shape changes dominates. Before differentiation, smaller transient protrusions are observed around the edges of the cells (Fig. 2A&B). During the early stages of differentiation, actin is organized in more wave-like flows which seem to propagate down and along developing processes (Fig. 2C&D) indicated by gradations of color in a rainbow-like progression along a process. Finally, during the late stage of differentiation, we instead see most of the dynamics localized not along the processes but instead perpendicular to the cell processes (Fig. 2C&F).

**Figure 2.**
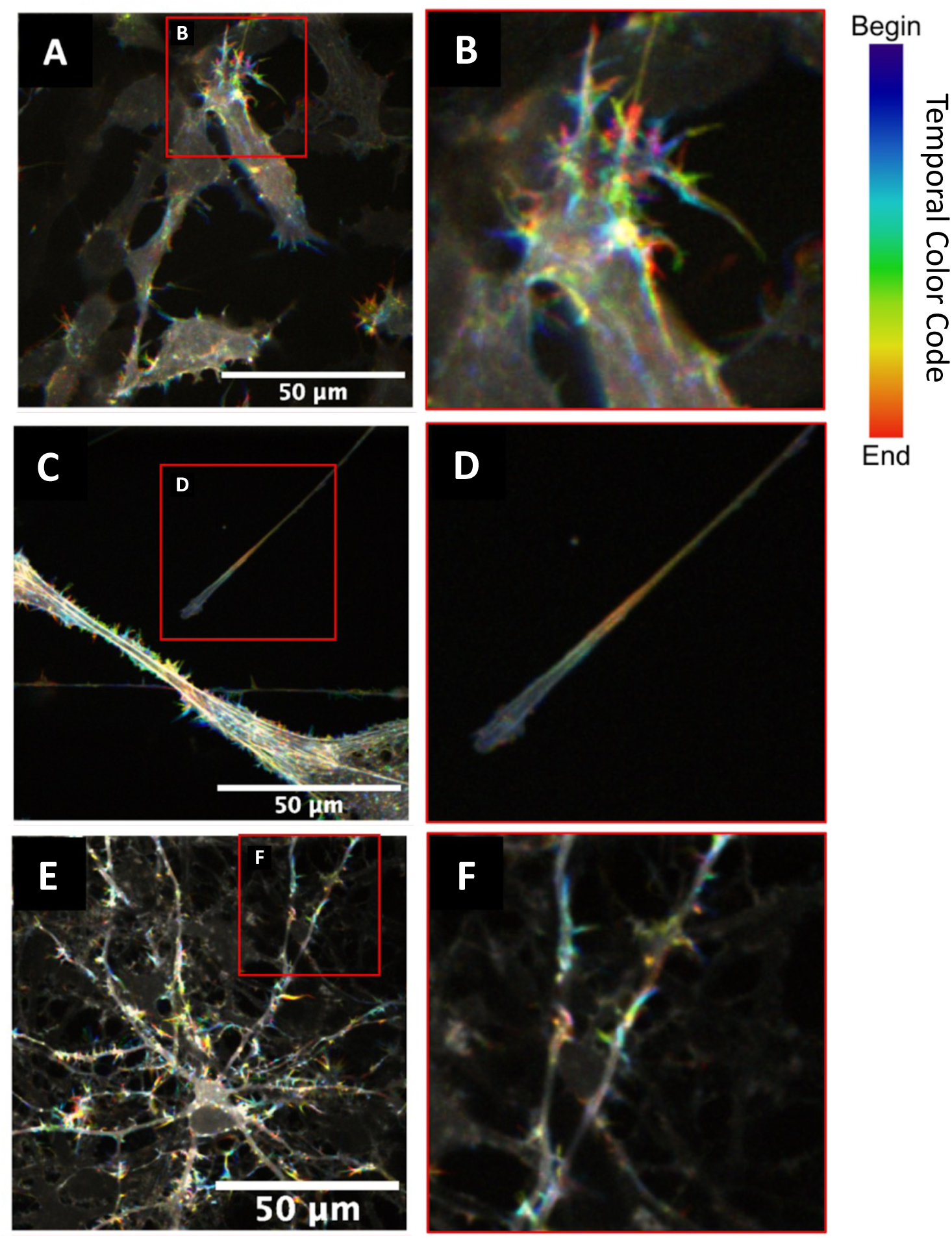
hNPC actin dynamics change scale over differentiation. Images represent temporal max projections. Areas that are white suggest static regions and areas of color show dynamics. hNPC in an undifferentiated state (**A**, zoom in on **B**). hNPC early differentiation, 5 days post differentiation initiation (**C**, zoom in on **D**). hNPC in mature differentiated cells, 21 days post differentiation initiation (**E**, zoom in on **F**).

To understand the directionality of actin flow throughout differentiation we employed optical flow (Fig. 3A-D) ^49–51^. Optical flow is a type of computer vision algorithm that compares subsequent frames of an image sequence and interprets the directionality and magnitude of flow based on the pixel. Representative images show a single frame from an undifferentiated cell using optical flow to analyze actin dynamics showing the optical flow vectors’ magnitude, orientation, and histogram of the speed of actin pixels in the single frame (Fig. 3A, B, and C respectively) as well as analysis of the entire film sequence showing the orientation of optical flow vectors over time (Fig. 3D). It is important to note the detected optical flow speed matches known speeds of actin polymerization waves, with the typical fast waves reported to travel approximately 1 micron per minute. Optical flow was implemented on undifferentiated, early differentiated, and mature differentiated cell time series (n = 16). The top 15% of magnitude flow vectors from each frame were retained and used to construct images showing where the highest optical flow was observed in each time series (Fig. 3E-G). Undifferentiated cells show broad areas of activity over time mainly within the cell body (Fig. 3E), early differentiated cells showed regions of activity along processes (Fig. 3F), while late differentiated cells show most activity among small punctate regions that appeared along and perpendicular to processes (Fig. 3G). This fits well with the generally accepted view that neuronal cells are maximally dynamic in their morphology during the early stages of differentiation when cell fate is determined. Propagation of actin waves would be expected to be seen during the projection of axon and dendrite growth cones away from the soma ^30,52^. Finally, smaller dendritic branches and synapses form after contact between processes, potentially leading to dynamics more like the observed punctate actin activity.

**Figure 3.**
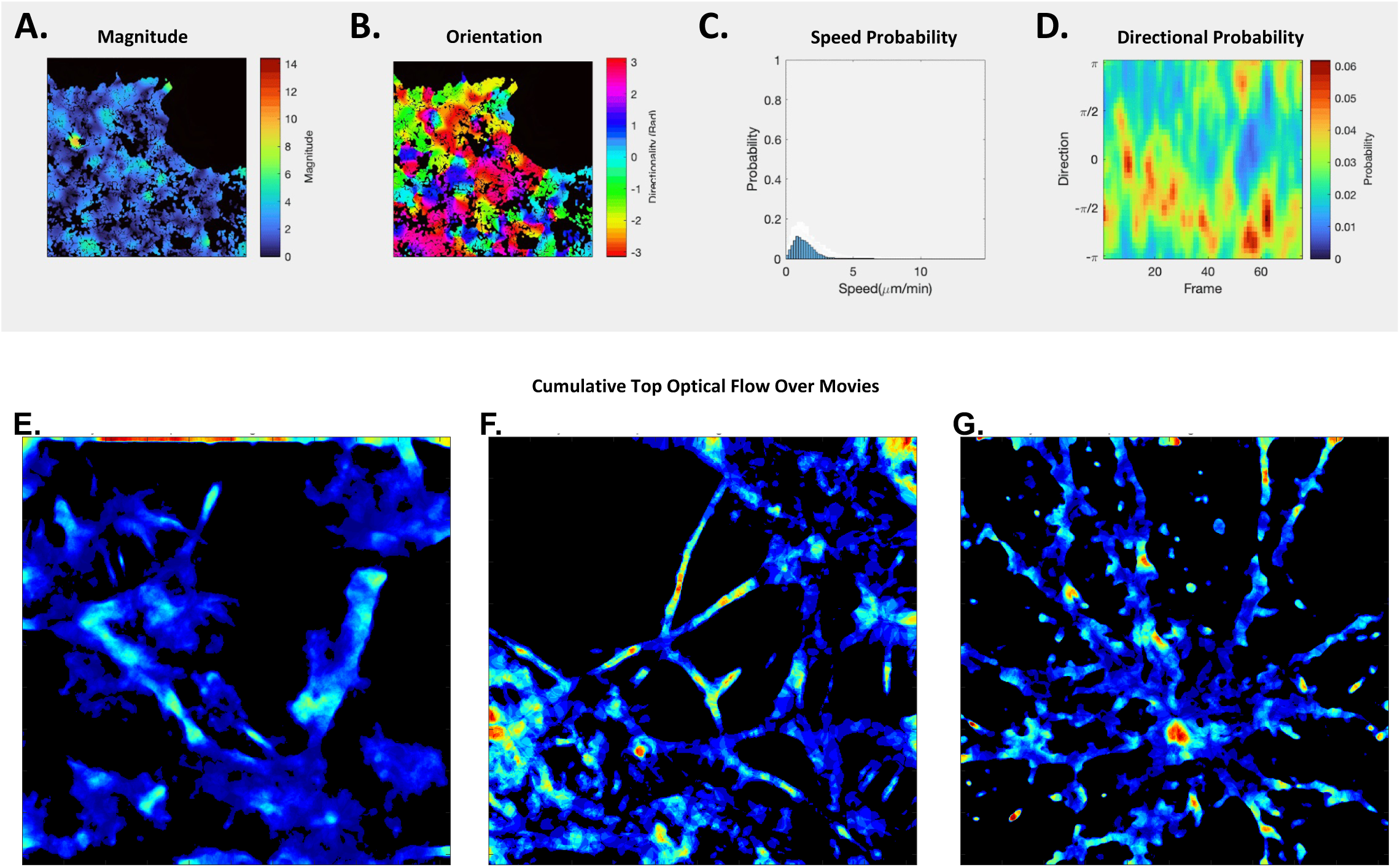
Optical flow as a tool to study actin dynamics with top 15% optical flow vectors show distinct regions of activity throughout differentiation. Actin dynamics within hNPC can be captured using optical flow algorithm (**A-D**). Single frame from a 160-second movie (acquired at 0.5 Hz) showing intracellular actin labeled (**A-C**). The magnitude calculated by optical flow algorithm is overlain cell region (**A**). Orientation of optical flow vectors displayed overlain cell region (**B**). Probability histogram of optical flow detected actin speeds in a single frame (**C**). Kymograph that captures the probability (as indicated by color) of directional actin flow (as indicated on the y-axis) over time in frames (along the x-axis) (**D**). Max projection of actin image time-series showing top 15% magnitude flow vectors. The “hotter” the color, the more flow vectors are found in multiple frames. In undifferentiated cells (**E**), early differentiated cells (**F**), and mature differentiated cells (**G**).

To further illustrate the changes in actin dynamics throughout differentiation, using optical flow, we analyzed actin flow directionality along the major axis of the cell over 5-minute intervals. Using optical flow, our assay is able to pick up the directionality of actin flow by analyzing actin dynamics from frame to frame to interpret the direction of an actin polymerization. When compiling the entire 5-minute image sequences, we found that actin flow is consistently polarized along the cell’s major axis throughout the different stages of differentiation (Fig. 4A-C). When looking over individual frames for the image sequences we see that cells consistently show actin flow along the cell’s major axis throughout time (Fig. 4D-F). Along with biased actin flow along the major axis, one key feature seen in the analyzed kymographs is that throughout hNPC development, actin dynamics appear to have a rhythmic character that is maintained throughout differentiation (Fig.4E-G).

**Figure 4.**
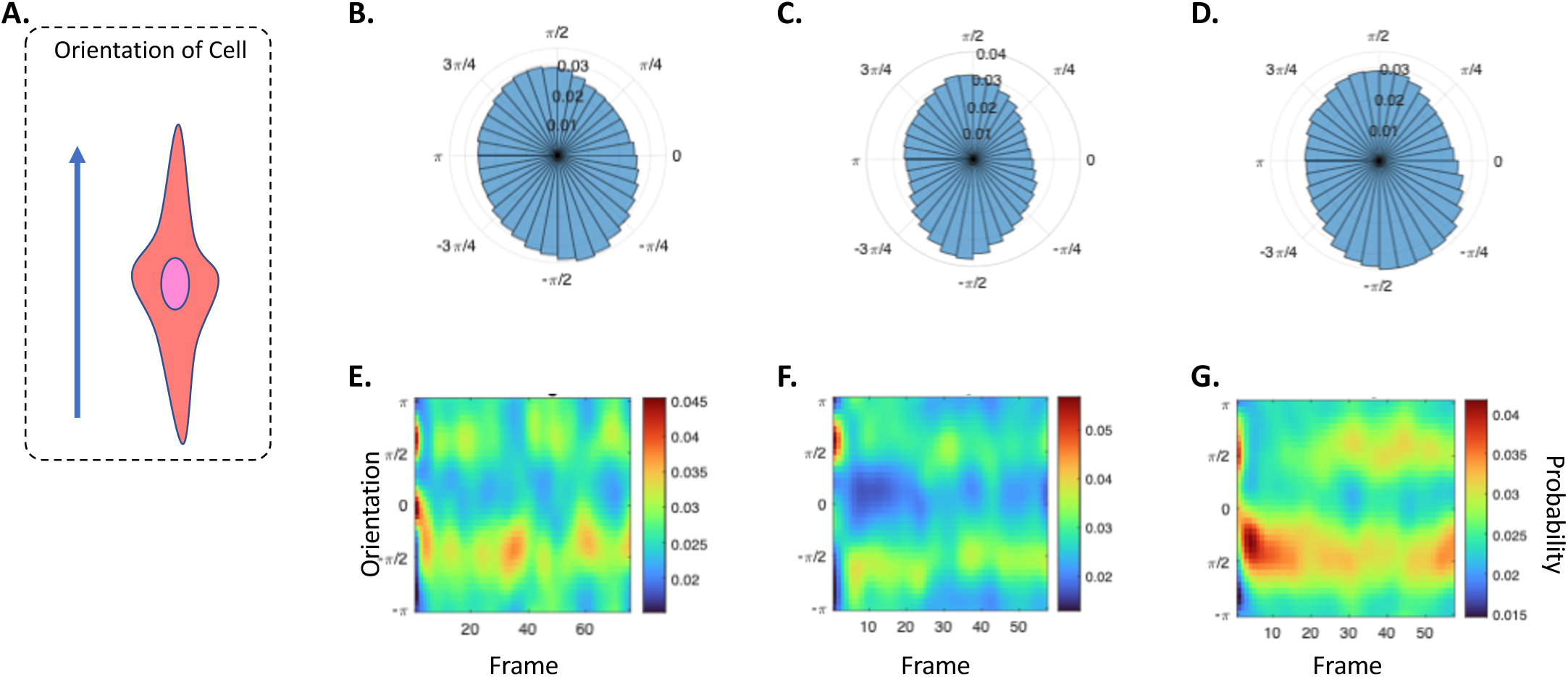
Actin dynamics have a characteristic rhythm and are oriented along the cell’s major axis throughout differentiation. Schematic with an arrow showing orientation along a cell’s major axis (**A**). Representative cell samples under various stages of differentiation: Undifferentiated (**B, E**), naïve differentiated (**C, F**), and mature differentiated (**D, G**). Rose plots (**B-D**) show optical flow vector orientation over entire 3-minute movies. Lower heatmaps (**E-G**) show orientation along the y-axis, time in frames along the x-axis, and probability as indicated by the color bar. Hotter colors indicate a higher probability. In all stages of differentiation looked at, there is a clear bias in optical flow vectors along the y-axis orientation.

### Automated Analysis and Representation of Population-Level Calcium Dynamics in hNPC

Calcium dynamics within developing hNPC are associated with proliferation, signaling, and developing cell-to-cell communication. To investigate the communication and coordination of calcium signaling throughout differentiation, calcium dynamic data from hNPCs are visualized using the cell-permeant dye Calbryte590, a calcium fluorescent dye that increases in intensity with increases in intracellular calcium concentration. Calbryte590, a non-ratiometric probe, was chosen due to its brightness, intracellular retention, and compatibility with the experimental protocols. Actin and calcium dynamics were not followed in parallel in any of these experiments since these dynamics are on different timescales (actin over tens of seconds, calcium over hundreds of milliseconds) and distinct length-scales (intracellular for actin vs cell scale for calcium) calling for separate analysis. Populations of about 100-300 cells in each field of view were imaged (Fig. 5A). This methodology and our automated analysis pipeline allowed us to look at individual cells and extract the fluorescent intensity over time. Individual cells can be identified either in an automated or manual fashion and labeled at the cell center (Fig. 5B). Fluorescent intensity over time is collected at 3Hz with 100ms exposure time (Fig. 5C) and represented in a kymograph (Fig. 5D). This frame rate is fast enough to yield multiple data points for each observed calcium transient. Using this automated workflow, the kymograph representation allows us to then plot all cells along the y-axis with time represented along the x-axis and look at the dynamic changes in calcium intensity over time for large populations of cells (Fig. 5E, F). The representative kymograph obtained from the analysis in figure 5 shows a mixture of active and inactive cells detected during analysis which is standard for our *in vitro* hNPC neural populations (Fig. 5). This disparity of active and inactive cells is seen in previous research, and our downstream analysis excludes non– or low spiking cells ^53^.

**Figure 5.**
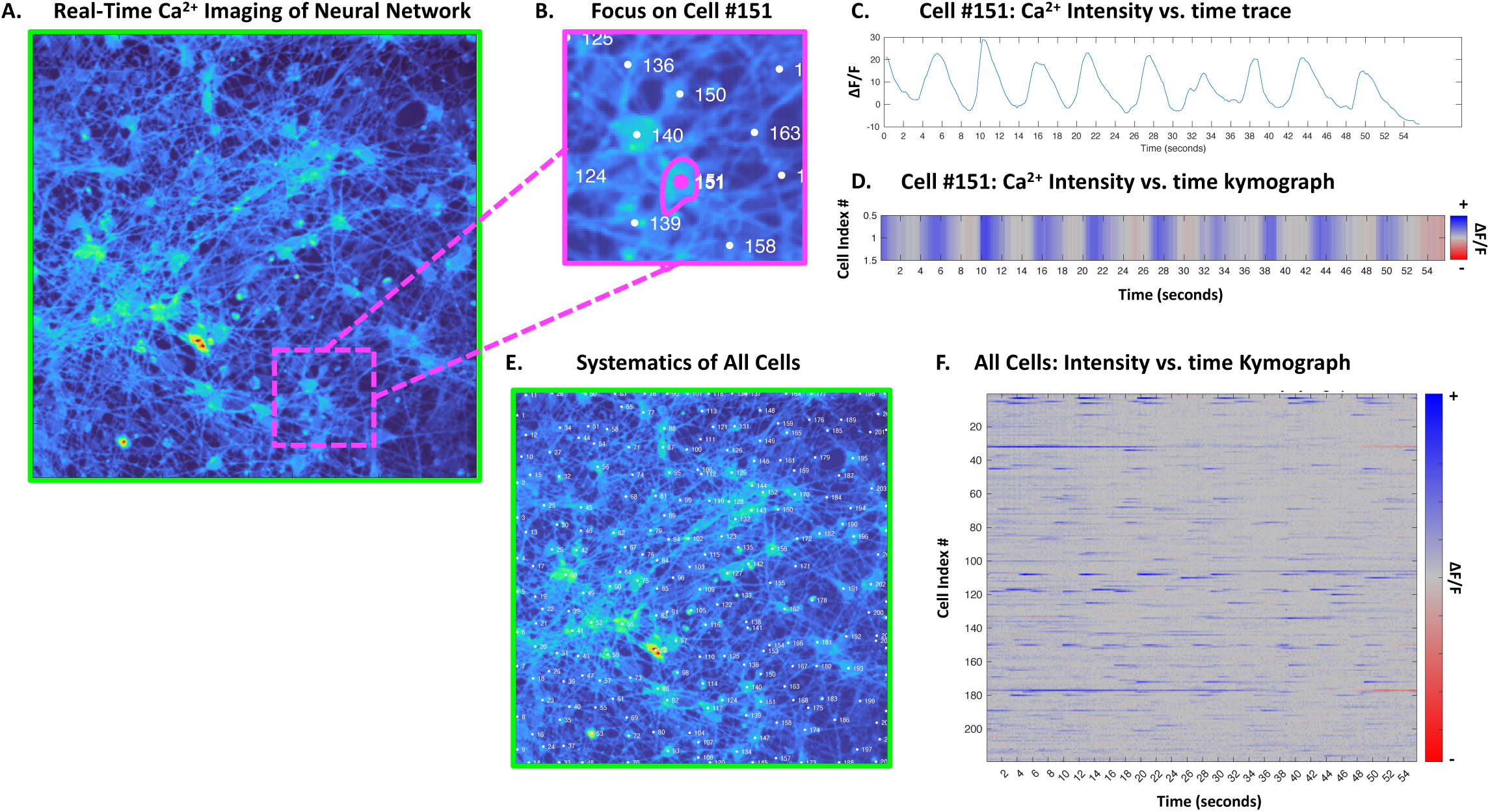
hNPC calcium dynamics can readily be quantified at the population level. Microscope field of view of hNPCs,10 days post differentiation initiation (**A**). Zoom in from A highlighting individual cell #151 in magenta (**B**). Trace of calcium fluorescent data over time from cell #151 (**C**). Kymograph representation of trace from C for cell #151; time on the x-axis, fluorescent intensity indicated as color (**D**). Automated cell centers are displayed as dots and numbered from the entire microscope field of view from A (**E**). Kymograph representing all cell fluorescent intensities over time (**F**).

### Pharmacological arrest of hNPC actin modulates Calcium Dynamics

Finally, to understand the impacts of actin dynamics on the calcium transient dynamics within developing hNPCs we use a pharmacological cocktail of drugs to inhibit actin dynamics during the naïve differentiation stage of our cells ie: day 4-5 after differentiation initiation (n = 13). The pharmacological cocktail of the ROCK inhibitor Y-27632, latrunculin A, and jasplakinolide (JLY cocktail) has been shown previously to arrest actin dynamics over a period in multiple cell types, while still allowing for other excitable dynamics like calcium signaling. To arrest dynamics initially the ROCK inhibitor is added for 10 minutes, before the addition of latrunculin A and jasplakinolide. For this study, we were able to show the arrest of actin dynamics in 4-5 days post differentiation initiation hNPCs using the JLY cocktail and correspondingly observed the impact of actin arrest on the ionic calcium dynamics within hNPCs (Fig. 6). In control conditions temporal color-coded max projections display regions of color suggesting activity over time (Fig. 6A) whereas cells treated with the JLY cocktail show stark black and white images (Fig. 6B) indicating static structures. These results suggest JLY cocktail addition works well in stabilizing actin structures and inhibiting dynamics. In control conditions using temporal color-coded max projections show calcium dynamics observed for a 1-minute interval show few colors suggesting less dynamic calcium activity (Fig. 6C) whereas the same interval of time in JLY, actin arrested conditions show many more colors suggesting an increase in activity (Fig. 6D). These changes are further quantified an illustrated in the population level kymograph representations of individual cell fluorescent intensities over time (Fig. 6E,F). Control conditions (Fig. 6E) show less activity compared to the JLY, actin arrested condition (Fig. 6F) as indicated by an increase in the number of blue peaks that occur representing activity in the kymographs. This is further quantified in the following section. In summary, actin dynamics within actin-arrested conditions show decreased activity as indicated by more white regions in the actin-arrest condition compared to the control condition (Fig. 6A&B). Calcium dynamics within actin-arrested conditions show more increased activity as indicated by more colored regions in the actin-arrest condition (Fig. 6C&D) and a potentially increased frequency of spikes within the kymograph compared to the control condition (Fig. 6E&F).

**Figure 6.**
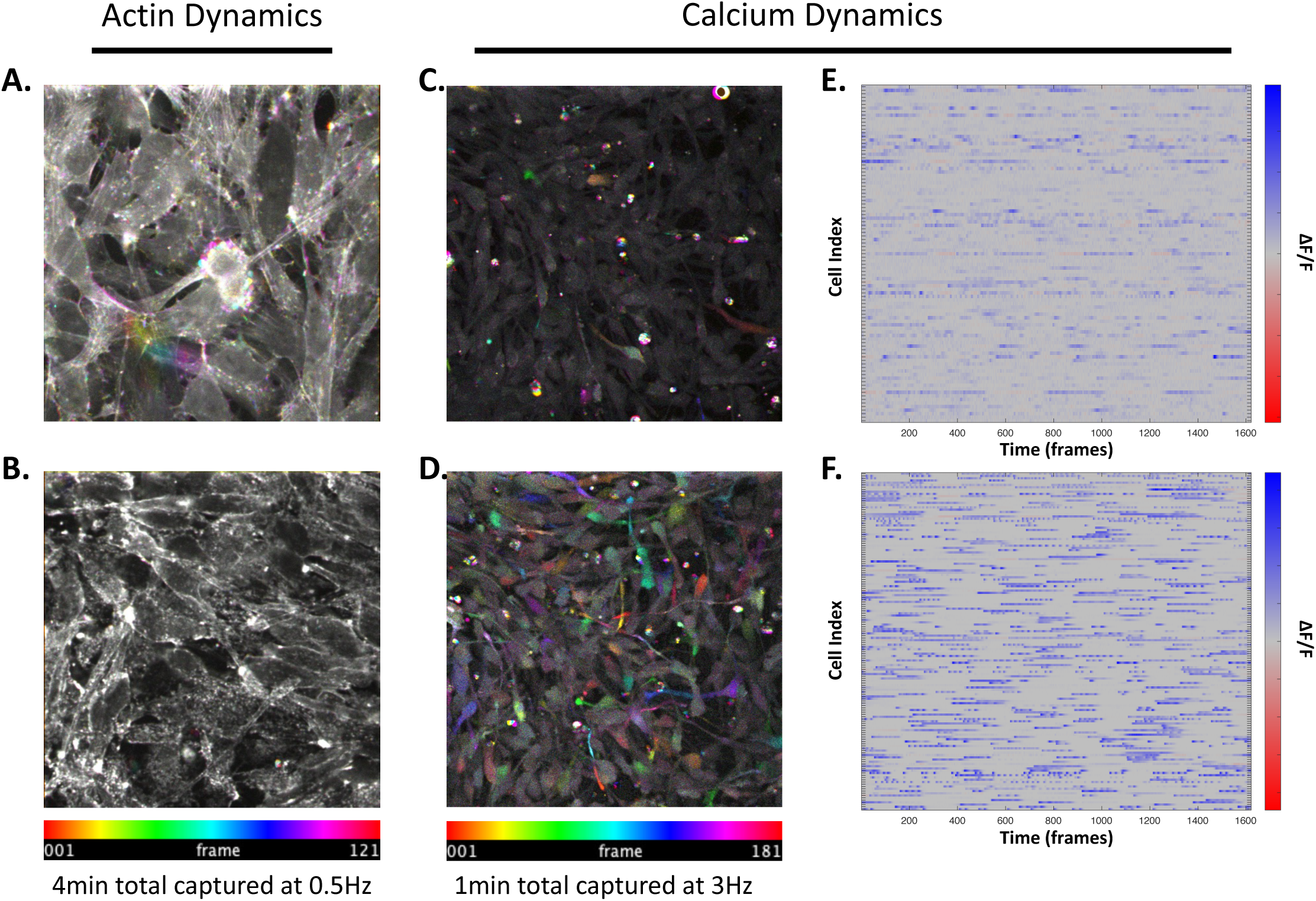
Pharmacological actin arrest in naive differentiated hNPC increases calcium dynamics. Temporally color-coded max projections from 4-5 days post differentiation initiation hNPCs of LifeactTagGFP2 over 4 min films captured at 0.5Hz during control or actin arrest (**A** & **B** respectively). Temporally color-coded max projections from Calbryte590 over 1 min films captured at 2Hz during control or actin arrest (**C** & **D** respectively). Kymographs showing calcium dynamics over 9 minutes captured at 2Hz with fluorescent intensity indicated by the color bar on the right-hand side, and individual cells indexed along the y-axis during control or actin-arrested conditions (**E** & **F** respectively).

### Pharmacological arrest of hNPC actin dynamics increases frequency of calcium transients

With the previous results observed we then further quantified the impact of actin arrest on the ionic calcium dynamics observed. What we found was that pharmacological inhibition of actin dynamics does not statistically increase the proportion of active cells, i.e. those with at least 3 transients as detected using a spike detection algorithm, within the population compared to the control condition (Fig. 7A). The proportion of active cells between control conditions in medium vs actin arrest in medium was not statistically significantly different from one another (t-test comparison p = 0.0620). Interestingly, by comparing the actin arrested condition to a control condition with excitable Tyrode’s solution, we found that cells in Tyrode’s had a significantly higher proportion of active cells compared to actin arrest in growth medium (t-test comparison p = 0.022874). Based on observations from the films as well as the changes seen in the kymographs of population activity (Fig. 6E&F) we then looked at quantifying the spiking frequency as measured as the number of spikes per population of active cells over the films’ time. We found there was a statistically significant increase in the calcium transient frequency of the population of active cells when we compared the JLY, actin arrest condition to the control condition (t-test comparison p = 0.010755) (Fig. 7B) in the cell growth medium. Furthermore, to test whether actin arrest leads cells to a maximally enhanced excited state, the medium was switched to the excitable Tyrode’s solution. With actin-arrest + Tyrode’s solution, we observed an even greater increase in the number of active cells with calcium dynamics within the population (Fig. 7A, green box) as well as an increase in the frequency of calcium dynamics (Fig. 7B, green box) which were found to be statistically significant (t-test comparison p = = 0.0024182 & p = 0. 026676 respectively) compared to actin arrest alone. We also note that cells without actin arrest in Tyrode’s solution compared to cells with actin arrest in Tyrode’s solution had significantly different proportions of active cells (Fig. 7A magenta and green boxes, t-test comparison p = 0.0062029) yet cells that were active had similar spiking frequencies (Fig. 7B magenta and green boxes, t-test comparison p = 0.36104) to one another. To better understand how this increase may be related to network activity, we further analyzed the data to obtain Pearson’s correlation coefficient values between all pairs of active cells. For the four conditions (control in medium, control in Tyrode’s, actin arrest in medium, and actin arrest in Tyrode’s solution) we then plotted the distribution of probability values from the range –1 to 1. What we found is that actin arrest leads to a narrowing of the distribution of correlation coefficient values compared to the control condition (Fig. 7C, in both the red curve and blue curve respectively), but that is very similar to Tyrode’s solution on control cells (Fig 7C, magenta curve). And that this narrowing of the correlation coefficients is not ameliorated in the case of Tyrode’s solution (Fig. 7C, green curve), although the proportion of active cells and frequency of activity is much higher in this condition (green box, Fig. 7A&B respectively). Interestingly, both conditions with actin arrest (both in growth medium and excitable Tyrode’s solution, Fig. 7C red and green curves) show an increase in the number of positive correlations (closer to 1) when compared with cells under control conditions with Tyrode’s (Fig. 7C magenta).

**Figure 7.**
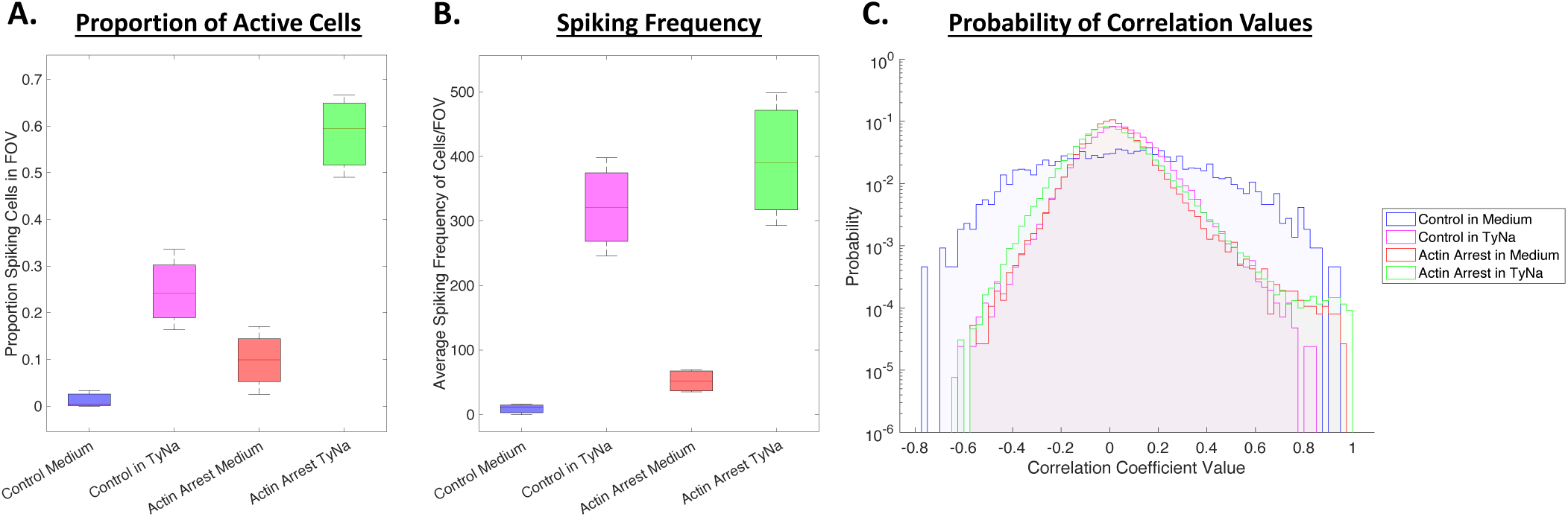
Actin arrest of naive differentiated hNPC does not impact the proportion of active cells but does change the frequency of active cells. Box plots showing the proportion of active cells under control condition in medium, control condition in excitable Tyrode’s solution, actin arrest in medium, and actin arrest in excitable Tyrode’s solution (**A**). Box plots showing the spiking frequency of active cells in populations under the same conditions as in A (**B**). Histograms showing the relative probability of correlation values from all active cells pairs under the same conditions as in A and B on a logarithmic y-axis (**C**).

## Discussion

This work focuses on the evolution of biomechanical activity during the development of neural networks, and includes two key aspects of neural networks that are missed in the typical neuron-centric and action potential-centric view of brain communication: that the composition of neural networks includes non-neuronal cells, and that neural cell excitability is not just electrical in nature. The contribution of this work is two-fold:

First, this study demonstrates that neural cells have reproducible actin dynamics and that the characteristic scale of actin dynamics changes with cell differentiation in a way that can be simply connected to changes in cell function. We show that actin dynamics change the spatial scale as a function of the differentiation stage – first being localized to emerging protrusions in undifferentiated cells, spanning axonal regions during early stages of differentiation, and becoming smaller, on synaptic scales, in late stages of differentiation. In addition to these well-known roles of actin dynamics at the leading edges of protrusions, growth cones, and nascent synapses, we find that the dynamics also occur away from such leading edges and that the dynamics have characteristics of an excitable system with rhythmic actin dynamics observed throughout differentiation ^31,32,38,54–56^. With this in mind, future work should look to further characterize the rhythmic actin dynamics within neural populations of cells, and to determine if the amplitude or frequency of that rhythm is associated with specific cell states or diseases. Future work should also seek to understand how modulating this rhythm may enhance neural cell differentiation, development, and communication.

Secondly, our study suggests that actin dynamics are not only driving the structural formation of the network but are an active participant in neural information flow. In hNPCs, differentiation leads to networks of functionally connected neural cells. Our readout of the collective character of the network is the collective calcium dynamics, which we show are impacted directly via modulation of structural actin dynamic arrest. It has been long established that cytoskeletal rearrangement is necessary and plays a key role in the maintenance of synaptic vesicles ^54,57^, and recent work has shown that dynamic restructuring of the cytoskeleton, primarily at dendritic spines and synapses, can play a key role in neural signaling ^29,32,58,59^. One study, in particular, found that mechanical stimulation of presynaptic boutons enhanced neurotransmission for greater than 20 minutes, through glutamate release and SNARE protein recruitment ^29^. This raises the question of whether intrinsic rhythmic intracellular actin dynamics play an active role in information transfer within a neural network. When we arrest the biomechanical rhythms, we observe that the frequency of calcium transients increases, and at the same time correlations decrease significantly. This result raises the possibility that more subtle modulations of actin dynamics may be harnessed to alter neural network activity and network link characteristics. These are two characteristics of networks that impact the information processing capability of the network. This work suggests that future experiments should look at how modulation of actin dynamics impacts the release of primary neural signaling molecules (neurotransmitters and gliotransmitters) and ion channel functioning. We note one potential caveat that must be understood is the impact of LifeActGFP2 on nascent intracellular actin dynamics. While LifeAct is commonly used to study live intracellular actin dynamics, it is known that the addition of this small protein probe can perturb actin (though less so than a direct fusion protein like actin-GFP)^48,60,61^. Another potential study limitation pertains to calcium buffering, which might be influenced by the presence of calcium indicators. These indicators, crucial for monitoring calcium dynamics, can subtly impact actual calcium concentrations in cells, potentially affecting result accuracy^62^. Nevertheless, our study aimed to compare calcium dynamics between actin-arrested and non-arrested cells, emphasizing the relevance of our findings in understanding cellular behavior within the context of actin regulation. This study only looked at mixed populations of neural cells, future work should look to distinguish the potential for modulation of actin in selectively isolated neurons or astrocytes to determine the impacts of changing actin rhythms in a cell-type specific manner has on the larger population and network activity dynamics.

Our discovery that actin dynamics impact neural network function raises the intriguing possibility that synaptic strengths themselves may experience rhythmic changes in connection strengths, thus giving the neural network a unique plastic characteristic where second to minute timescale fluctuations in link strengths coexist with fast ionic information processing. Indeed, this slow biomechanical rhythm may provide an intrinsic plasticity that gives living neural networks their unique ability to adapt quickly and to store and transmit information on a wide range of timescales. Previous work has already suggested the biomimetic inclusion of systems analogous to astrocytes can be used to improve ANN ^24,63,64^. Thus, as an important next step, our work suggests it may be important to assess the consequences of rhythmic plasticity for the performance of biomimetic ANN algorithms.

## Methods

### Cell culture

hNPC with v-myc transgene were cultured in accordance with previous methods. Briefly, cells were expanded on Matrigel-coated plates or tissue culture flasks (NEST and Corning). Cells were maintained in growth medium composed of DMEM:F12+GlutaMax (Thermo Fischer Scientific) supplemented with 2% B-27 plus neural cell supplement (Thermo Fischer Scientific), 1% penicillin-streptomycin (Sigma-Aldrich), [50.]mM heparin (Stemcell Technologies), in the presence of 10 ng/mL bFGF (Stemcell Technologies) and 20 ng/ml EFG (Stemcell Technologies). Differentiation medium was composed of the same components as growth medium, excluding the growth factors (bFGF and EGF). Cells were maintained in a cell culture incubator at 37°C in a humidified environment with 5% CO_2_. Cells were passaged at around 95% confluency. Briefly, cells were rinsed with DPBS (brand), and cells were detached using a brief incubation with accutase (Stemcell Technologies) for 5-10 minutes one volume of DMEM:F12+GlutaMax was added. The cells are centrifuged at 500 g for 5 minutes. The cell pellet was resuspended in fresh growth medium before being plated on Matrigel-coated plates. All experiments were carried out on cells between passages 5 and 30.

### Generation of LifeactGFP cell line

hNPC line was cultured as described above. rLV-Ubi-LifeAct-TagGFP2 lentivirus purchased from Ibidi technologies (catalog no. 60141) was used according to supplier instructions, with the addition of viral transduction reagent on cell cultures around 50% confluent at desired MOI (2 and 4). After 48 hours of incubation for viral uptake, media containing viral particles was replaced. Cell samples were then checked under a confocal fluorescence microscope for the presence of a GFP signal. Cells were subjected to antibiotic selection using 5 ug/mL puromycin to establish stable polyclonal lines. After 2-7 days of selection, cells were expanded with samples cryopreserved for future use. Lifeact-generated lines were used between passages 7 and 30.

### Calcium Labeling and Imaging

Intracellular calcium labeling was prepared using Calbryte-590 AM (AAT Bioquest) following manufacturer-provided protocol. A stock solution is prepared in anhydrous DMSO. Briefly, appropriate dilution of Calbryte-590 AM was added to cell medium to achieve 1uM final working concentration. Cells were then incubated for 30-60 minutes in a 37°C incubator. The cell sample was then incubated at room temperature for another 15 minutes. Afterward, the cell sample medium was replaced with an appropriate imaging solution (either cell culture growth medium, Tyrode’s solution, or Tyrode’s solution with supplemented sodium). Samples were then immediately imaged on a system with temperature, humidity, and CO2-controlled environments using 561 nm lasers to excite fluorescent indicators.

### Pharmacological arrest of actin

The pharmacological arrest of actin was performed according to previous literature using modified concentrations.^65^ For the JLY treatment, first, hNPC are incubated with 10 μM Y-27632 for 10 minutes. Next jasplakinolide (3 μM) and latrunculin B (5 μM) were added. For calcium imaging with actin arrest, cells are preincubated with Calbryte-590 AM for 30-60 minutes as described above before having the medium replaced, after which Y-27632 is added for 10 minutes before the addition of jasplakinolide and latrunculin B.

### Immunocytochemistry

At specific endpoints cell samples were removed from the incubator, the medium was removed, and cells were rinsed with PBS for 5 minutes. Cell samples were then fixed in 4% paraformaldehyde/PBS for 15 minutes, followed by rinsing three times in PBS for 5 minutes each. Cells were then permeabilized with 0.1% Triton x 100 + 1% BSA in PBS for 1 hour at room temperature. Direct immunofluorescence was then performed using specific antibodies incubated overnight at room temperature. Nestin was probed using anti-nestin-AF-488 at 5:1000 (Santa Cruz Biotechnology, catalog no. sc-23927 AF546, 10c2 clone). Sox2 was probed using anti-sox2-AF-546 at 5:1000 (Santa Cruz Biotechnology, catalog no. sc-365823 AF546, E-4 clone). βIII-tubulin was probed using anti-TUBB3-AF-640 at 5:1000 (BD Pharmingen, catalog no. AB_1645400, TUJ1 clone). Glial fibrillary acidic protein was probed using anti-GFAP-AF-546 at 2:1000 (Santa Cruz Biotechnology, catalog no. sc-58766 AF546, GA5 clone). After direct antibody staining overnight, the antibody solution was removed, and cells were rinsed in PBS for 5 minutes before being stained with Hoechst (1:2000) in PBS for 2 to 4 minutes followed by an additional PBS wash.

### Imaging cell cultures

Image sequences of fixed cell cultures for immunocytochemistry and live imaging of actin and calcium dynamics were captured on a PerkinElmer spinning disk microscope using 405 nm, 488 nm, 561 nm, and 640 nm wavelength lasers at 10% power. Live cultures were maintained using a Tokai Hit environmental chamber with temperature, CO2, and humidity control.

### Image Analysis

All image analysis conducted using either the ImageJ processing package FIJI or using Matlab. FIJI (ImageJ) used to obtain temporal color-coded max projection images. First Image sequences from the microscope opened using ImageJ and converted into 8-bit. Next, the temporal color code plugin by Kota Miura applied. On images of actin dynamics, jitter is removed using the image registration plugin “Linear Stack Alignment with SIFT using. Looking at images, most drift or jitter was identified as translational motion, therefore the expected transformation option was selected to be translational. In noisy images, the FIJI plugin “remove outliers” was used with a radius of 2 pixels to remove bright outliers.

To analyze actin dynamics, In Matlab, optical flow was implemented using the “estimateflow” function using the “<u>opticalFlowFarneback</u>” object with the following parameters: the number of pyramid layers 7, filter size 10, neighborhood size 1, pyramid scale 0.5. Optical flow vectors from samples from various stages of differentiation were then used to create the plots in figure 3 and figure 4.

In Matlab, Tiff image sequences are loaded in. Cell centers are then identified (automatically using a difference of gaussian approach and then identifying cell centers as local maxima) of an expected cell radius size and used to record ROI of fluorescent activity over time. Fluorescent activity for each cell is normalized by dividing the raw fluorescent intensity by the maximum value. Moving average baseline subtraction is performed to compute the delta F over F (ΔF/F). These ΔF/F traces for each cell correspond to plots in Figures 5E, 5D, 5C, 6C, and 6E. Calcium spikes are detected using the “findpeaks” function with the stipulations for the minimum prominence value of 20 of the normalized ΔF/F value and maximum peak width of either 15 or 30 seconds. The number of peaks found was then used to calculate the proportion of active cells (with more than 3 peaks) under specific conditions as well as the frequency of spiking activity under specific conditions. For statistical data analysis of the calcium spike data, the Matlab “Ttest2” was implemented in a pairwise fashion amongst all pairs of the four final conditions. Results are shown in in Figure 7.

## Data Availability

The raw fluorescent microscopy recording data, analyzed films, and data files supporting the findings of this study are available upon reasonable request to the corresponding authors.

## Code Availability

FIJI was used to visualize fluorescent data images shown in figure 1, 2, & 6. The source code used for analysis in this study is available for public access on GitHub at the following link: https://github.com/SJG3/Coupled-Biomechanical-and-Ionic-Excitability-in-Developing-Neural-Cell-Networks_Code-Availability.

OF_lite_Farneback_SJG_availability.m was used to generate analysis shown in figure 3 & 4. DFF_code_availability.m was used to generate analysis shown in figure 5 & 6. Spike_detection_availability.m was used to generate analysis used in figure 7. We are committed to transparency and reproducibility in our research, and we encourage interested researchers to visit the GitHub repository for access to the source code used. For any additional inquiries or assistance, please contact the corresponding author Wolfgang Losert at.

## Acknowledgements

We thank the imaging core at University of Maryland College Park for their technical support with the equipment used. We would also like to thank all members of the Losert lab for their helpful discussions and feedback at various points during this research.

